# Comparative genomics sheds light on niche differentiation and the evolutionary history of comammox *Nitrospira*

**DOI:** 10.1101/138586

**Authors:** Alejandro Palomo, Anders G Pedersen, S Jane Fowler, Arnaud Dechesne, Thomas Sicheritz-Pontén, Barth F Smets

**Affiliations:** Department of Environmental Engineering, Technical University of Denmark, Kgs Lyngby, Denmark; Department of Bio and Health Informatics, Technical University of Denmark, Kgs Lyngby, Denmark

## Abstract

The description of comammox *Nitrospira* spp., performing complete ammonium-to-nitrate oxidation, and their co-occurrence with canonical betaproteobacterial ammonium oxidizing bacteria (β-AOB) in the environment, call into question the metabolic potential of comammox *Nitrospira* and the evolutionary history of their ammonium oxidation pathway. We report four new comammox *Nitrospira* genomes, constituting two novel species, and the first comparative genomic analysis on comammox *Nitrospira*.

Comammox *Nitrospira* has lost the potential to use external nitrite as energy and nitrogen source: compared to strictly nitrite oxidizing *Nitrospira*; they lack genes for assimilative nitrite reduction and reverse electron transport from nitrite. By contrast, compared to other *Nitrospira*, their ammonium oxidizer physiology is exemplified by genes for ammonium and urea transporters and copper homeostasis and the lack of cyanate hydratase genes. Two comammox clades are different in their ammonium uptake systems. Contrary to β-AOB, comammox *Nitrospira* genomes have single copies of the two central ammonium oxidation pathway genes, lack genes involved in nitric oxide reduction, and encode genes that would allow efficient growth at low oxygen concentrations. Hence, comammox *Nitrospira* seems attuned to oligotrophy and hypoxia compared to β-AOB.

β-AOBs are the clear origin of the ammonium oxidation pathway in comammox *Nitrospira*: reconciliation analysis indicates two separate early *amoA* gene transfer events from β-AOB to an ancestor of comammox *Nitrospira*, followed by clade specific losses. For *haoA*, one early transfer from β-AOB to comammox *Nitrospira* is predicted – followed by intra-clade transfers. We postulate that the absence of comammox genes in most *Nitrospira* genomes is the result of subsequent loss.

**Significance:** The recent discovery of comammox bacteria - members of the *Nitrospira* genus able to fully oxidize ammonia to nitrate - upset the long-held conviction that nitrification is a two-step process. It also opened key questions on the ecological and evolutionary relations of these bacteria with other nitrifying prokaryotes. Here, we report the first comparative genomic analysis of comammox *Nitrospira* and related nitrifiers. Ammonium oxidation genes in comammox *Nitrospira* had a surprisingly complex evolution, originating from ancient transfer from the phylogenetically distantly related ammonia-oxidizing betaproteobacteria, followed by within-lineage transfers and losses. The resulting comammox genomes are uniquely adapted to ammonia oxidation in nutrient-limited and low-oxygen environments and appear to have lost the genetic potential to grow by nitrite oxidation alone.

## Introduction

Nitrification, the biological oxidation of ammonium to nitrate, is an essential process in terrestrial and aquatic environments and is essential in water quality engineering applications. For the last century, nitrification was assumed a two-step process executed by two complementary functional groups, ammonia-oxidizing prokaryotes (AOP) and nitrite-oxidizing bacteria (NOB). Recently, several groups have shown that single microorganisms belonging to the genus *Nitrospira* can carry out the complete oxidation of ammonia to nitrate, a process abbreviated *comammox* (1–4). *Nitrospira* spp., known as strict nitrite-oxidizers, are widespread in both natural and engineered ecosystems associated with nitrogen cycling (5, 6). The *Nitrospira* genus is extremely diverse and comprises at least six lineages, which frequently coexist (5, 7). Comammox *Nitrospira* genomes described to date belong to *Nitrospira* lineage II, and comprise two clades (clade A and B) based on the phylogeny of their ammonia monooxygenase (1, 2). Both clades were detected in samples retrieved from a groundwater well and in a groundwater-treating rapid sand filter with conventional *Nitrospira* spp (1, 4). The diversity of *Nitrospira* spp. points towards ecological niche-partitioning; nitrite and dissolved oxygen concentrations are potential niche determinants for *Nitrospira* lineages I and II (8, 9). The recently described metabolic versatility of some *Nitrospira* spp., which includes formate, hydrogen, urea and cyanate metabolisms (10–12) may also contribute to their coexistence. Nevertheless, little is known about differences in the functional potential of the two comammox *Nitrospira* clades and niche-partitioning between comammox *Nitrospira* and AOP. Only the hypothesis that comammox *Nitrospira* could outcompete canonical AOB in surface-attached and substrate-limited environments has been suggested (13). The evolutionary history of ammonia oxidation in *Nitrospira* is unknown. Based on their ammonium monooxygenase (AMO) and hydroxylamine dehydrogenase (HAO) sequences, b-proteobacterial ammonium oxidizing bacteria (AOB) are the most closely related to comammox *Nitrospira* (1–4). Either AMO and HAO encoding genes are ancestral in *Nitrospira* spp. or were acquired by horizontal gene transfer from AOP, even though the degree of divergence makes a recent acquisition improbable (2).

To examine the functional potential which might niche-partitioning among comammox *Nitrospira* and between comammox *Nitrospira* and AOP, and to unravel the evolutionary history of ammonia oxidation in *Nitrospira*, we performed differential coverage genome binning on a *Nitrospira* composite population genome recovered from a metagenome in which comammox *Nitrospira*, canonical AOB and ammonium oxidizing archaea (AOA) co-occurred (4, 5). Comparative genomics analysis was conducted with the recovered genomes and high quality published *Nitrospira* genomes. The evolutionary history of genes involved in ammonia oxidation in comammox *Nitrospira* was also explored with a special focus on understanding the potential role of horizontal gene transfer, gene duplication, and gene loss. For this purpose, we executed pairwise protein dissimilarity comparison between and within comammox clades as well as with other AOB, explored the genomic arrangement in the relevant pathways, and performed probabilistic and parsimony-based reconciliation analysis where trees for individual genes were compared to a clock-like species-tree constructed from ribosomal proteins. Our study revealed comammox-, comammox clade-and canonical *Nitrospira*-specific features related to nitrogen assimilation, electron donor versatility and substrate-limitation tolerance. Our analysis strongly suggests that genes for the ammonium pathway were acquired by transfer from b-proteobacterial ammonia oxidizers to comammox *Nitrospira*, and occurrence of additional transfer events (for some AO related genes) between comammox clade A and clade B.

## Results and Discussion

### Genome extraction from the metagenome

Metagenomes from adjacent locations in a RSF were used for individual genome extraction (4). Five *Nitrospira* genomes (CG24_A, CG24_B, CG24_C, CG24_D, CG24_E) were recovered from the metagenomes using differential coverage (14) and a subsampling assembly strategy (Fig. 1). The characteristics of these and other genomes used in this study are shown in Table 1. The size of the comammox genomes assembled here range from 3.0-3.6Mb, with a completeness of 95-100%. Based on phylogenetic analysis of 14 syntenic ribosomal proteins, all genomes belong to *Nitrospira* Lineage II (Fig. S2). Genes required for complete ammonium oxidation (*amo* and *hao* operons) were detected in four of the genomes. Based on *amoA* phylogeny CG24_A, CG24_C and CG24_E belong to comammox clade B and CG24_B to comammox clade A (Fig. S3). CG24_A was most abundant at an average relative abundance of 18.6% (of all metagenome sequence reads) followed by CG24_B (8.6%), CG24_C (4.6%), CG24_D (2.4%, the only canonical NOB *Nitrospira* recovered) and CG24_E (1.1%). (Fig. S4).

**Figure 1.**
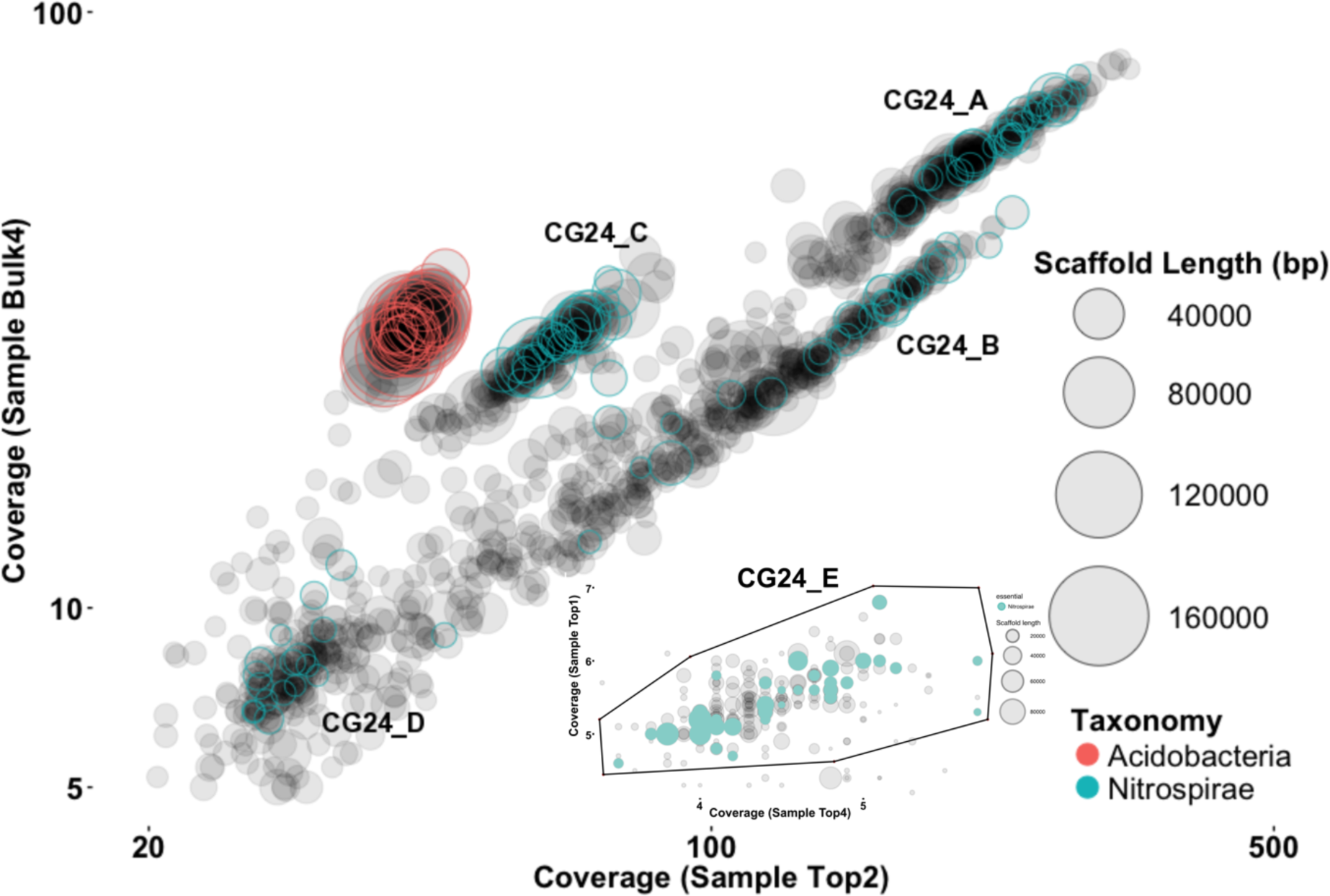
Differential coverage plot of two metagenomes obtained from Islevbro waterworks at different sampling depth. Scaffolds are displayed as circles, scaled by length and colored according to phylum-level taxonomic affiliation. Only scaffolds >4 kbp are shown. A second differential coverage plot showing the CG24_E bin extraction (enclosed by the polygon) is presented.

**Table 1.**
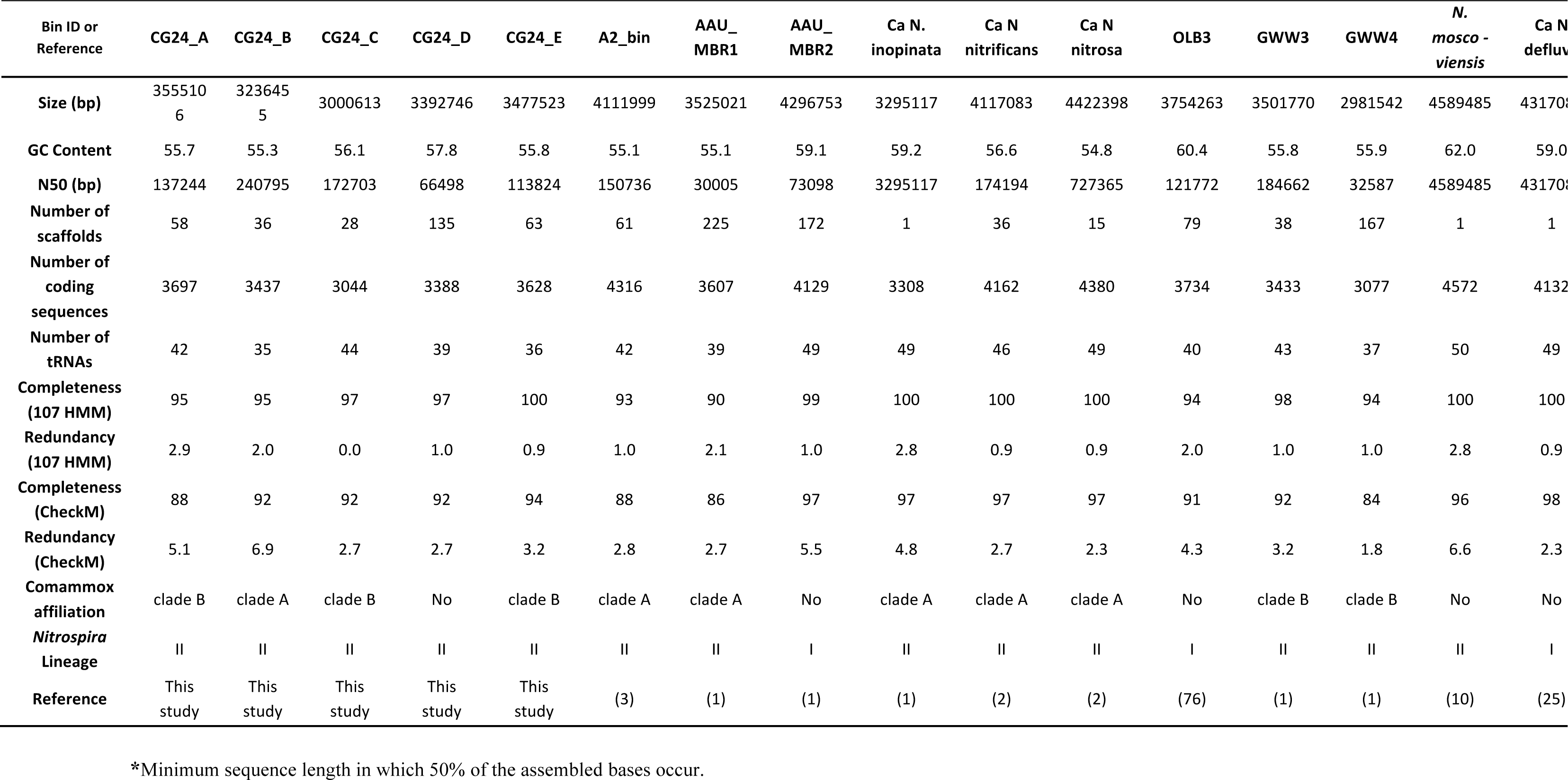
Characteristics of examined genomes.

### Comparative genomics

The newly recovered genomes were compared with each other and publicly-available, high quality (>90% completion, <5% redundancy) *Nitrospira* genomes, including 11 comammox and five canonical *Nitrospira* (Table 1). The 16 studied *Nitrospira* genomes constitute 11 different species, seven of them comammox *Nitrospira*. Based on average amino acid identity (AAI) analysis CG24_A, CG24_D and CG24_E are divergent enough from each other and previously published genomes to be separate species (at species level cut-off of 85% AAI similarity (15)) (Fig. S5).

The coding sequences (CDS) of the 16 genomes were annotated and grouped into SEED subsystems(16). Of the 59,744 CDS, 33.6% (range: 30% to 40%) were functionally annotated by SEED subsystems. *Nitrospira* lineage I genomes clustered together and were characterized by an overrepresentation of genes related to carbohydrate and aromatic compound metabolism, and synthesis of cofactors/vitamins (PCA analysis, Fig. S6). *Nitrospira moscoviensis*, although classified as lineage II, shared characteristics with lineage I genomes, including an overrepresentation of genes associated with virulence and defence genes. In contrast, the comammox *Nitrospira* genomes grouped separately from the previously described strictly nitrite oxidizing *Nitrospira*. Most of these genomes are enriched in genes related to amino acid metabolism, cell cycle and division, cell wall and capsule biosynthesis, and RNA metabolism.

More detailed comparison of the *Nitrospira* genomes was based on pangenomic analysis (Fig. 2). The 59,744 CDS of the 16 *Nitrospira* genomes clustered into 12,337 protein clusters (PCs), with a core *Nitrospira* genome consisting of 1,382 PCs. The core genome includes genes for the nitrite oxidation pathway, the reductive tricarboxylic acid cycle for CO2 fixation (rTCA), and the oxidative TCA cycle. Chlorite dismutase and copper-containing nitrite reductase (*nirK*) are also present in the core genome. 35 comammox-specific PCs were identified; 16 and 3 of these PCs had highest sequence similarity to homologs in β-proteobacterial ammonium oxidizing bacteria (β-AOB) and methane oxidizers, respectively (Table S2). In addition, we detected 57 and 52 comammox clade A and clade B-specific PCs, (Table S2). The specific metabolic characteristics identified in the pangenomic analysis are described below, summarized in Fig. 3 and the distribution of all mentioned genes in the examined genomes can be found in Table S2.

**Figure 2.**
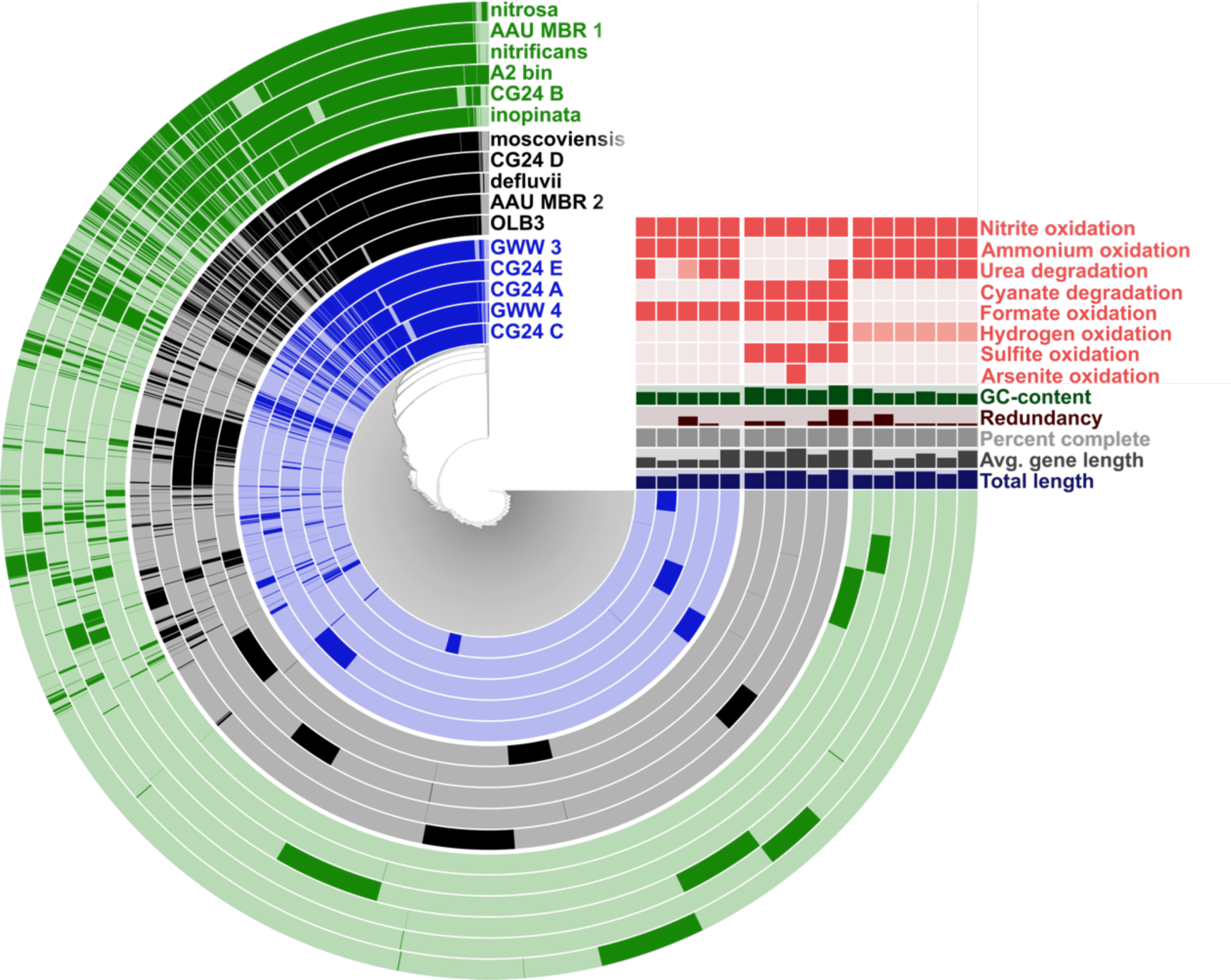
Communalities and uniqueness in the *Nitrospira* pangenome as derived from the clustering of 16 genomes based on 12,337 protein clusters (PCs). Each radial layer represents a genome, and each bar in a layer represents the occurrence of a PC (dark presence, light absence). Clade A comammox *Nitrospira* genomes are denoted in green and Clade B in blue.

**Figure 3.**
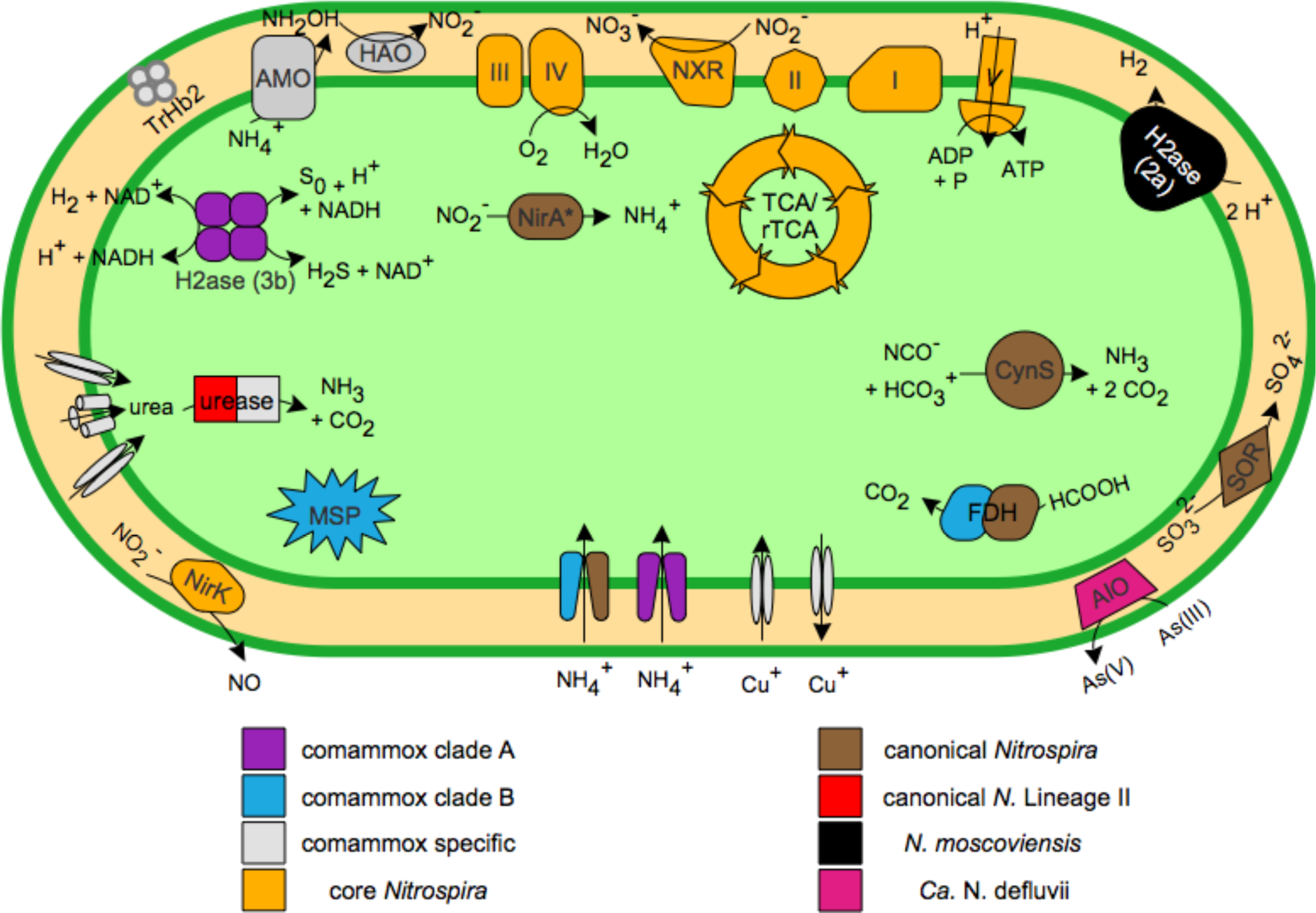
Cell cartoon of the core and specific key metabolic features in *Nitrospira* pangenome, as predicted from genome annotation. AIO, arsenite oxidase; CynS, cyanate hydratase; FDH, formate dehydrogenase; H2ase, hydrogenase; MSP, methionine salvage pathway; SOR, sulfite dehydrogenase; TrHb2, 2/2 hemoglobin group 2. Enzyme complexes of the electron transport chain are labeled by Roman numerals. * *N. moscoviensis* possesses a octaheme cytochrome c (OCC) putatively involved in nitrite reduction to ammonia.

### Nitrogen metabolism

All the recovered genomes encoded the nitrite oxidoreductase (NXR), consistent with other *Nitrospira* spp. The copy number of NXR in the investigated genomes varied from one, in most of the comammox genomes, to five in *N. moscoviensis* (Table S2). The ammonium oxidation pathway (AMO structural genes *amoCAB* and the putative AMO subunits *amoEDD2*) is present in four of the newly recovered genomes (CG24_A, CG24_B, CG24_C and CG24_E).

In terms of nitrogen uptake, a low-affinity Rh-type ammonium transporter with sequence similarity to homologs in β-AOB is only present in clade A genomes, while a high-affinity AmtB-type transporter was detected in clade B and canonical *Nitrospira* genomes. Difference in ammonium uptake affinity might, therefore, be an important niche-separation factor between the comammox genomes. Besides the uptake of exogenous ammonium by *Nitrospira*, ammonium can be produced through urea degradation as most of the comammox genomes harbour urease genes. This enzyme, which is functional in *N. moscoviensis* (10), is not detected in genomes of Lineage I *Nitrospira*. The comammox genomes contain a diversity of urea transporters. In addition to the high affinity urea ABC transporters (*urtABCDE*) present in some *Nitrospira* spp. such as *N. lenta* (10) (*N. moscoviensis* contains only *urtA*), comammox genomes also harbour two additional urea transporters: an outer-membrane porin (*fmdC*) involved in uptake of short-chain amides and urea at extremely low concentrations (17), and an urea carboxylase-related transporter (*uctT*). Moreover, comammox genomes uniquely harbour an additional agmatinase gene which is phylogenetically distinct from the one typically present in *Nitrospira* spp. and closely related to those found in *Actinobacteria* spp. This enzyme hydrolyzes agmatine, producing putrescine and urea. Taken together, this suggests that comammox *Nitrospira* may have a competitive advantage in urea uptake with respect to other *Nitrospira* in environments with low and/or fluctuating urea concentrations. On the other hand, cyanate hydratase genes (*cynS*) are only detected in canonical *Nitrospira*. This may confer a benefit over comammox in environments with very low ammonium concentrations and the presence of cyanate coupled with reciprocal feeding with ammonium oxidizers (12). All genomes except *Ca.* N. inopinata encode a 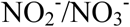 transporter gene (*narK)*. Furthermore, canonical *Nitrospira* genomes contain additional transporters for the uptake of nitrite: lineage I *Nitrospira* and CG24_D genomes harboured a nitrite transporter (*nirC)* and a formate/nitrite family transporter, while *N. moscoviensis* and CG24_D contained a nitrite-nitrate ABC transporter (*nrtABC*). The comammox genomes lack an assimilatory nitrite reductase (*nirA*) or octaheme cytochrome c (OCC) (which Koch *et al*. (10), proposed to be associated with nitrite reduction to ammonia in *N. moscoviensis*), which would prevent comammox *Nitrospira* growth in the presence of nitrite as sole N source, as observed in *Ca*. N. inopinata (1).

### Alternative electron donors

Some *Nitrospira* can utilize substrates beyond nitrite as electron donors including formate and hydrogen (10–12). The genes coding for formate dehydrogenase (*fdh*) are present in canonical *Nitrospira* and clade B comammox but not in clade A comammox. Thus, this provides an opportunity for niche-differentiation particularly in oxic–anoxic transition zones where formate is commonly found as a product of fermentation. The genes for the hydrogen producing enzyme, formate hydrogenlyase (FHL), were found only in canonical *Nitrospira* genomes as well as three comammox clade A genomes (A2_bin, AAU_MBR_1 and *Ca*. N. nitrosa). Although this protein is not functional in some *Nitrospira* spp. (11) it is not known if this is a common feature for all the genomes containing FHL reported here. With respect to hydrogen oxidation, the group 2a [NiFe] hydrogenase and accessory proteins involved in aerobic hydrogen oxidation in *N. moscoviensis* (11) are absent from the other *Nitrospira* genomes. However, all comammox clade A genomes encode a complete group 3b [NiFe]-hydrogenase. Although this bidirectional hydrogenase (sulfhydrogenase) is extensively distributed across the bacterial domain (18), little is known about its actual role. For the NOB, *Nitrospina gracilis,* Lücker *et al*. (19) proposed that sulfhydrogenase may play a role in the oxidation of H2 to provide reduced NAD(P)H for biosynthesis or energy generation, or in the reduction of polysulfide either for sulfur assimilation or as an electron acceptor during fermentation. Further studies are needed to unravel the function of this protein and their role in clade A comammox *Nitrospira*. Comammox genomes do not contain genes for a periplasmic sulfite:cytochrome c oxidoreductase (sulfite dehydrogenase, *sorAB*), which is characteristic of canonical *Nitrospira*, and would suggest their capacity for sulphite oxidation, given their genetic inventory for sulfur assimilation, as described for another NOB (*Nitrospina gracilis*) (19). Lastly, a complete arsenite oxidase was exclusively found in *Ca.* N. defluvii. Although all examined genomes harbour arsenic resistance genes, only *Ca*. N. defluvii has the genetic potential for energy generation during arsenite oxidation (SI Results and Discussion).

### Energy conservation and transduction

All *Nitrospira* genomes harbour two homologous sets of genes encoding the complex I of the electron transport chain (NADH:ubiquinone oxidoreductase, NUO). One of these NUO lacks the subunits involved in NADH binding and oxidation. This feature has been observed in other prokaryotes (20); in *Nitrospira* spp. this incomplete complex I may be used for reverse electron transport from quinol to low-potential ferredoxin, as proposed in *Nitrospina gracilis*(19), thereby providing reduced ferredoxin required for CO_2_ fixation in the rTCA cycle. Additionally, CG24_A, *Ca*. N. nitrosa, AAU_MBR1, A2_bin and *N. moscoviensis* possess a third complete homologous set of genes coding for complex I, which is phylogenetically divergent from the other two, and is closely related to the gammaproteobacterial clade E type. This type of complex I has also been found in other distantly related organisms, including *Nitrosospira multiformis* and some *Bacteroidetes* spp., suggesting acquisition by horizontal gene transfer (21). Although the exact function of this additional complex I is not known, the differential expression of distinct complex I isozymes has been observed under different conditions, pointing towards physiological versatility (22, 23). Also, differences with respect to complex II, the succinate:quinone oxidoreductase (SQR), were noted across genomes. SQR connects the TCA cycle to the quinone pool. This complex is classified in five types based on the number of membrane-bound domains and the presence or absence of a heme b (24). *Nitrospira* genomes do not all contain the same SQR type. Clade B, CG24_D, and genomes belonging to *Nitrospira* lineage I contain a SQR type E, whereas clade A genomes and *Nitrospira moscoviensis* possess a SQR type B. Differences were also detected for the cytochrome *bc* complex (complex III) which is involved in the transfer of electrons from quinol to a *c*-type cytochrome. The canonical *Nitrospira* genomes contain at least two copies of complex III (three in *N. moscoviensis* and OLB3), while the comammox genomes harbour just one. The absence of a second homologue of complex III might indicate that these comammox *Nitrospira* are not able to direct electrons from nitrite oxidation to reverse electron transport, impeding growth on nitrite alone as confirmed for *Ca*. N. inopinata (1). With respect to cytochrome oxidases (complex IV), the studied genomes harbour four subunits similar to the functionally transcribed ‘cyt *bd*-like oxidase’ (NIDE0901) of *Ca*. N. defluvii associated with nitrite oxidation (25). In addition to this terminal oxidase, canonical *Nitrospira* (except CG24_D), comammox clade A genomes (excluding *Ca*. N. nitrosa) and the clade B GWW3 bin contain the genes *cydAB* encoding a heterodimeric cyt. *bd* quinol oxidase, which could be related to the oxidation of alternative electron donors.

### Carbon metabolism

Comammox *Nitrospira*, like other sequenced *Nitrospira* genomes (25), encode genes for glycolysis, the pentose phosphate pathway and the TCA cycle. Moreover, clade B comammox genomes and AAU_MBR2 possess an acetate permease (*actP*). *Nitrospira* genomes have the potential to degrade catechol and protocatechuate (Table S2). Although these degradation pathways are only partially complete in *Nitrospira* genomes, comammox *Nitrospira* possess an intradiol ring-cleavage dioxygenase, which could transform catecholate derivatives to TCA cycle intermediates (26). Additionally, clade B comammox and *N. moscoviensis* harbour a 4-oxalocrotonate tautomerase, essential in the conversion pathway of various aromatic compounds such as catechol to intermediates for the TCA cycle(27). The *phaZ* gene which codes for polyhydroxybutyrate (PHB) depolymerase involved in PHB degradation was exclusively found in *Nitrospira* comammox genomes. The absence of this enzyme in other Nitrospirae species and the high sequence similarity with *Methylococcaceae* spp. suggest that comammox genomes could have acquired this feature through horizontal gene transfer from methanotrophs. Genes related to PHB synthesis were not found in the studied genomes, hence whether the PBH polymerase is a relic or has any functional role remains unknown. Overall, the presence of enzymes and pathways involved in different carbon sources degradation suggests that comammox *Nitrospira* has the potential to grow mixotrophically as reported for other *Nitrospira* spp. (28–30).

### Stress response, resistance and defence

Genes encoding catalase (*cat*) or superoxide dismutase (*sod*), providing protection against reactive oxygen species (ROS), are absent from the closely related comammox genomes *Ca*. N. nitrosa and AAU_MBR1 bin as well as from *Ca*. N. defluvii. On the other hand, CG24_A and *N. moscoviensis* harbour diverse genes related to ROS protection including one catalase and one peroxidase (two in both cases for *N. moscoviensis*), and two dissimilar superoxide dismutases. The remaining genomes encode for either catalases or superoxide dismutases and all genomes harbour the putative ROS defence mechanisms predicted for *Ca*. N. defluvii (consisting of cyt. *c* peroxidases, thioredoxin-dependent peroxiredoxins, manganeses, bacterioferritin and carotenoids) (25). Contrary to strict nitrite-oxidizing *Nitrospira*, the comammox genomes contain a 2/2 hemoglobin type II (TrHb2), which has been associated with oxidative stress resistance and oxygen scavenging (31, 32). Thus, the distinct capacity to deal with oxidative stress within *Nitrospira* spp. may enable these organisms to coexist by occupying different microniches in environments with oxygen gradients such as biofilms and flocs. Most of the genomes encode two proteins homologous to RsbUV, related to environmental stress response, and clade A comammox genomes possess a fusion protein phylogenetically distinct from other *Nitrospira* spp. that is homologous to RsbUVW. *N. moscoviensis* also encodes all the genes for the stressosome (*rsbRSTX*), a complex that controls several signaling pathways in response to diverse environmental stresses (33). Although this stressosome was first characterized to be associated with the activation of sigma factor B, which is not present in gram negatives such as *Nitrospira* spp., other downstream stressosome-related pathways involving a putative CheY-like or a diguanylate cyclase have been described for *Vibrio brasiliensis* and *Moorella thermoacetica*, respectively (34, 35). *N. moscoviensis* possesses two genes homologous to CheY-like and diguanylate cyclase in the flanking regions of the stressosome cluster. Hence, based on these genomes characteristics, clade A comammox bacteria and canonical *N. moscoviensis* might be better adapted to changing environmental conditions than their counterparts. On another note, genes associated with low-resource environments were encountered in some of the *Nitrospira* genomes. Unlike canonical *Nitrospira*, the comammox genomes contain Cu^2+^ homeostasis genes (*copCD* and *copAB*) with highest sequence similarities to homologs in broteobacterial ammonium oxidizers. These proteins usually confer higher copper tolerance and increased Cu^2+^ uptake (36, 37), so could provide an advantage to comammox *Nitrospira* in environments with low copper availability. Canonical *Nitrospira* genomes harbour the genes for the cytochrome *c* biogenesis system II. In contrast, comammox genomes contain the genes for the cytochrome *c* biogenesis system I. While system II requires less energy for cytochrome synthesis, system I is advantageous in iron-limited environments as this system has hemes with higher iron affinity (38). As nitrifiers have a high iron requirement for their [FeS] cluster-and heme-containing enzymes, this could provide an important competitive advantage for comammox in iron-limited environments (39). Clade B comammox *Nitrospira* genomes contain the methionine salvage pathway which was not detected in clade A or canonical *Nitrospira* genomes (Fig. 3 and Table S2). This pathway is involved in the recycling of sulfur-containing metabolites to methionine and is upregulated under sulfur-limiting conditions (40, 41). Thus, clade B comammox could better persist during periods of sulfur depletion. Genes related to arsenic and mercury resistance were found in all of the studied genomes. Especially striking were the cases of comammox CG24_B and AAU_MBR1 which contain phylogenetically distinct chromate, arsenic and mercury resistance proteins in a putative integrative conjugative element (SI Results and Discussion).

### Comparison between comammox *Nitrospira* and ammonia oxidizing microorganisms

β-AOB genomes are characterized by the presence of two or three copies of *amo* and *hao* gene clusters, together with the presence of NOx detoxification pathways. These characteristics are considered to form the basis of the ecological success of this group in ammonia-rich habitats. In contrast, comammox *Nitrospira* generally harbour a single copy of *amo* and *hao* genes, like gammaproteobacterial AOB (g-AOB) and AOA, and lack the membrane-bound cytochrome *c* nitric oxide reductase (cNOR), heme-copper nitric oxide reductase (sNOR), nitrosocyanin, and cytochrome P460 (*cytL* gene), all usually present in AOB. This feature of comammox is shared with AOA, however the investigated comammox *Nitrospira* genomes contain two proteins related to resistance to nitrosative stress, a NO-responsive regulator (*nnrS*) and a 2/2 hemoglobin type I (TrHb1), that were not detected in AOA. Taken together, comammox *Nitrospira* appear adapted to environments with low ammonium concentrations and thus could be more directly in competition with oligotrophic AOP. The different AOP groups also clearly differ in their carbon fixation pathway. AOB and AOA utilize the oxygen-tolerant Calvin–Benson–Bassham and hydroxypropionate– hydroxybutyrate cycles, respectively, while comammox *Nitrospira* possess the microaerophilic related rTCA pathway. Furthermore, canonical AOP genomes encode the low-affinity *aa3*-type heme-copper oxidase (*Nitrosomonas eutropha* also contain a high-affinity cytochrome *c* oxidase *cbb3*). In contrast, comammox *Nitrospira* harbour high affinity cytochrome *bd*-like oxidases. Additionally, the 2/2 hemoglobin type II (TrHb2), associated with oxidative stress resistance and oxygen scavenging, which was detected in comammox *Nitrospira,* is not universal for AOB (only present in *Nitrosospira* spp.) and has not been detected in AOA. These three features together suggest that comammox *Nitrospira* could have a competitive advantage over AOB and AOA in microaerophilic environments. Another characteristic of comammox *Nitrospira* is its potential to compete under phosphorous and copper limiting conditions. Comammox *Nitrospira* genomes contain an alkaline phosphatase (*phoD*), which is highly expressed under phosphorus limitation and starvation (42, 43). This enzyme was not detected in AOA genomes and is not universal in AOB (putative homologs are present in *Nitrosomonas sp*. Is79A3*, Nitrosococcus halophilus* and *Nitrosospira* spp.). In relation to copper homeostasis, while *copCD* genes are present in both AOB and comammox genomes, the genes *copAB* detected in comammox *Nitrospira* are not common for AOB as just a few species seem to harbour homologues of these genes (*Nitrosomonas europaea, Nitrosomonas eutropha, Nitrosospira multiformis and Nitrosospira* sp. Nv17). Although AOA seem to have a high Cu^2+^ demand due to their high presence of copper-containing proteins, their Cu^2+^ acquisition mechanisms are unknown (44). These two aspects could be of great importance as both phosphorous and copper deficiency can impact nitrification in engineered environments (45–47). Taken together, comammox *Nitrospira* have the features of a slow growing organism adapted to nutrient-limited conditions. Consistently, comammox *Nitrospira* has so far mainly been detected in substrate-limited environments as well as in low oxygen environments and/or with microaerophilic niches (1, 2, 4, 48–50).

### Horizontal gene transfer events shaped comammox *Nitrospira*

The AMO and HAO sequences of β-AOB are most closely related to those of comammox *Nitrospira*(1–4). In fact, the sequence similarity of *amo* and *hao* genes between β-AOB and comammox *Nitrospira* (ca. 60% amino acid sequence identity for *amoA* and ca. 66% for *haoA*) is higher than between β-AOB and β-AOB (ca. 45% for *amoA* and ca. 53% for *haoA*), even though the latter two groups belong to the same phylum (Proteobacteria), and are distantly related to the Nitrospirae phylum. This apparent incongruence suggests horizontal gene transfer (HGT) of *amo* and *hao* genes between comammox and β-AOB. To explore whether HGT has played a role in the evolutionary history of comammox *Nitrospira* and β-AOB, we compared the sequence dissimilarity of proteins related to the ammonium oxidation pathway and 18 housekeeping proteins to that of a set of 14 ribosomal proteins (Table S1). We included sequences from the 11 comammox genomes investigated in this study as well as from eight previously published β-AOB genomes (Table S3). Pairwise dissimilarity comparisons showed that housekeeping proteins are linearly related to the ribosomal proteins across essentially all genomes (Fig. 4 and Fig. S7) confirming vertical inheritance. For most of the proteins associated with ammonium oxidation, the relationship was different (Fig. 4 and Fig. S8) with obvious discontinuities. In some cases, the sequence dissimilarities between β-AOB and comammox genomes were as high as comammox clade A and clade B were to each other, although these two groups are closely related based on ribosomal proteins (Fig. 4 and Fig. S8). Hence, the evolutionary history of these genes has been more complex, and supports the hypothesis of the occurrence of horizontal transfer(s).

**Figure 4.**
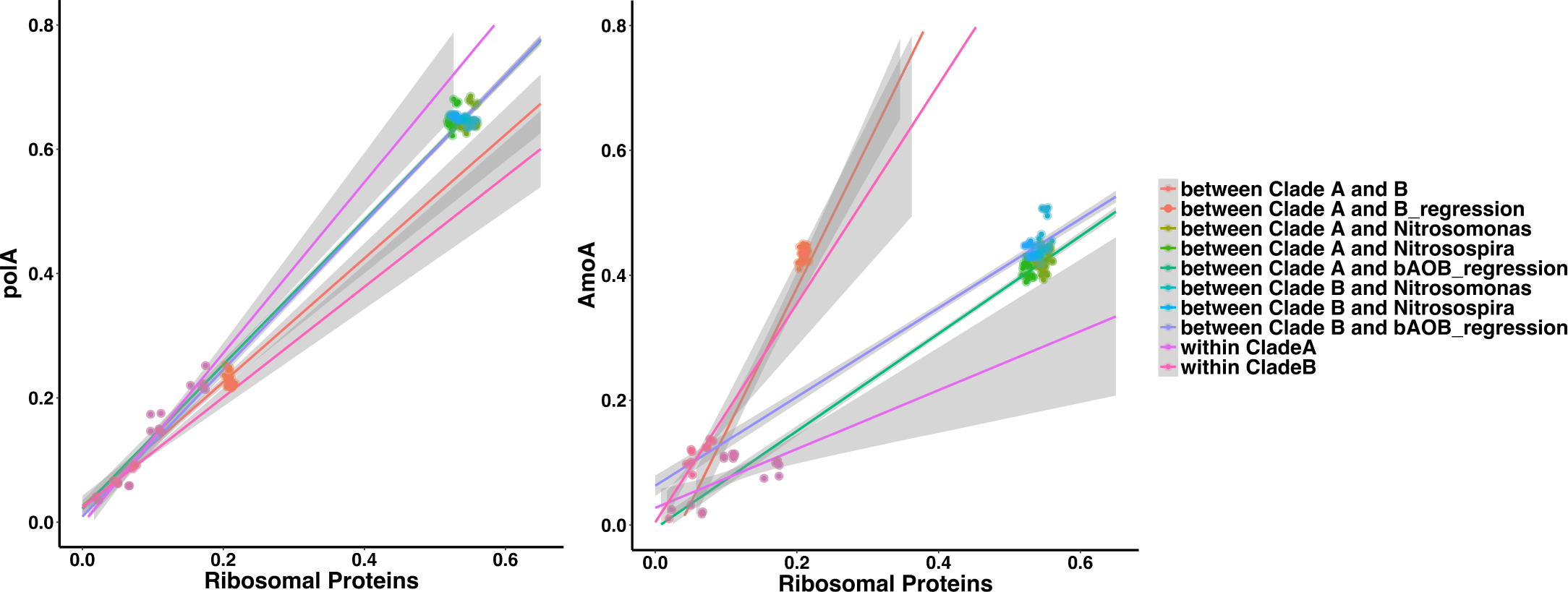
Relationship between genomes phylogenetic distance and protein sequence divergence for a house keeping protein (left) and for the ammonium monooxygenase (right) for comammox *Nitrospira* and β-AOB genomes. Each dot represents a pair of genomes and is coloured according to the groups to which the compared genomes belong. The y-axis shows the pairwise protein dissimilarity (fraction of amino-acid sites that differ) while the x-axis shows the corresponding pairwise dissimilarity for a set of 14 ribosomal proteins. Coloured lines show the linear regression for each groupwise comparison with shadowed regions indicating 95% confidence intervals for the slopes. A: comammox clade A; B: comammox clade B; Ntsm: *Nitrosomonas* genomes; Ntrssp: *Nitrosospira* genomes. AOB: Both *Nitrosomonas* and *Nitrosospira*.

We, subsequently, analysed the genomic arrangement of ammonium oxidation-related genes for all comammox genomes and for representative β-AOB genomes (Fig. 5 and Fig. S9). For all comammox bacteria except for *Ca*. N. inopinata, the AMO and HAO gene clusters as well as the cytochrome *c*-biosynthesis genes are situated in the same genomic region. This is not the case for AOB. Additionally, comammox bacteria have two copies of *amoD,* while AOB only have one copy. Among the comammox bacteria, clade B genomes uniquely contain a duplication of one of the cytochrome *c*-biosynthesis proteins (CcmI). Two duplicated genes associated with iron storage (*brf*) are found in clade A genomes next to the putative AMO subunits *amoEDD2* (SI Results and Discussion).

**Figure 5.**
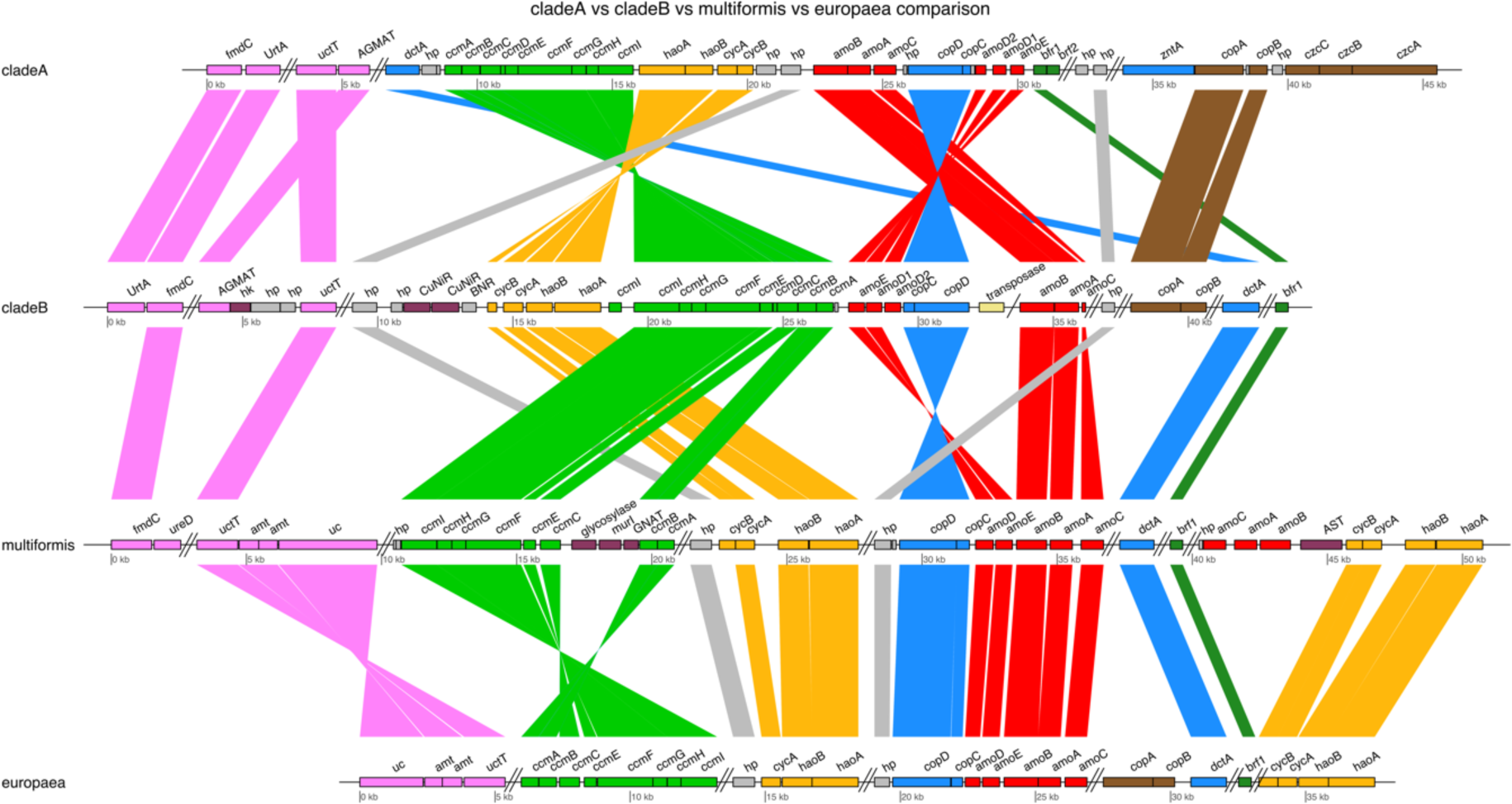
Schematic illustration of the ammonium oxidation pathway genomic region as well as other AOB-related genomic features in comammox *Nitrospira* clade A (*Ca.* N. nitrificans), clade B (GWW3 bin) and selected ammonia-oxidizing bacteria (*Nitrosospira multiformis* and *Nitrosomonas europaea)*. Homologous genes are connected by lines. Functions of the encoded proteins are represented by colour. Parallel double lines designate a break in locus organization. Single line designates a break probably due to contig fragmentation.

A reconciliation analysis was performed to stringently examine the possible occurrences of gene duplication(s), gene loss(es), and horizontal gene transfer(s). Here, we constructed a clock model-based species-tree from the 14 ribosomal proteins (Table S1), including the 16 examined *Nitrospira* genomes, (*Leptospirillum ferrooxidans*), eight β-AOB and two γ-AOB publically available genomes (Fig. S10 and Table S3). Gene trees were constructed for the ammonium oxidation-related genes under investigation (Table S3). The probabilistic (Bayesian) analysis provided strong support for horizontal transfer of the majority of the investigated genes (Table S4): for 12 genes the posterior probability for at least one transfer event was 95%-100%, for another 10 for the posterior probability was 80%-95%. For all investigated genes, we found very strong support (95%-100%) for either a transfer or a duplication event.

The evolutionary history of *amoA* and *haoA* was further investigated. For *amoA*, the reconciliation model suggests two separate early gene transfer events from β-AOB to an ancestor of comammox *Nitrospira*, followed by clade specific losses such that the genes in clade A and B originate from two separate HGT events (Fig. 6). Hence, for *amoA* the split between the two clades is much deeper than suggested by the species tree, which could explain the high dissimilarity between the *amoA* sequences of the otherwise closely related clade A and clade B comammox genomes (Fig. 4). In addition, a transfer of *amoA* from the common ancestor of *Ca.* N. nitrosa and AAU_MBR_1 to *Ca.* N. inopinata was inferred. This transfer would explain the deviation in the tetranucleotide pattern of the *amoCAB*-containing region compared to the genome-wide signature identified in *Ca.* N. inopinata by Daims *et al*. (1). Furthermore, the gene order in the *amo* operon in *Ca.* N. inopinata is different from the other comammox genomes, namely *amoCAB, hp, copDC* instead of *amoBAC, copDC* (Fig. S9). Unlike the other comammox genomes, *Ca.* N. inopinata does not have the *amo* operon located in the same genomic region as the *hao* operon and the genes for cytochrome c-biogenesis system II (Fig. S9).

**Figure 6.**
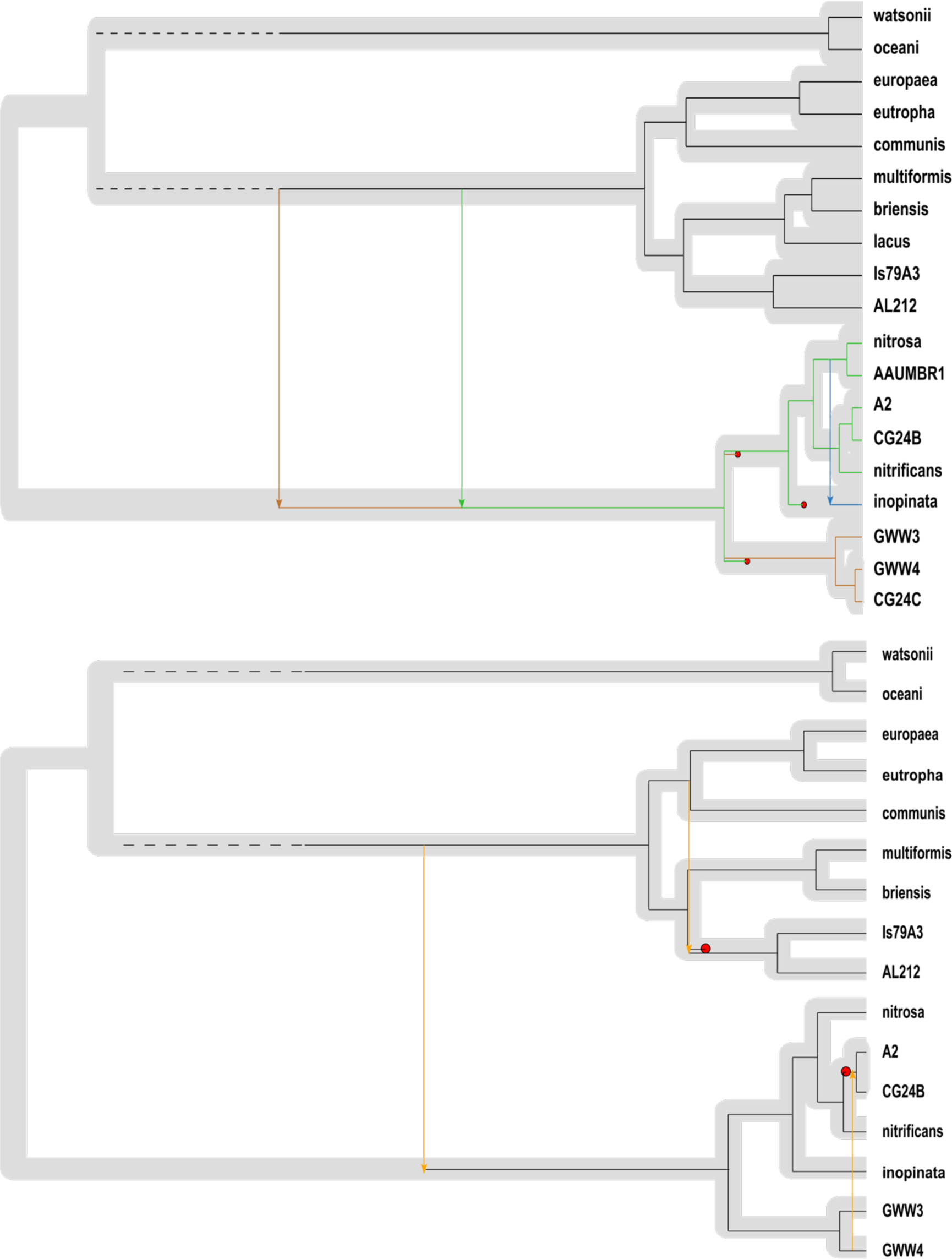
Reconciliation of functional gene trees (top. *amoA*; bottom. *haoA*) with species tree for comammox *Nitrospira* and β-AOB genomes. The species tree, based on 14 ribosomal genes, is shown in grey with the gene-trees super-imposed on top in narrower black lines. Arrows and red dots denote transfer and loss events, respectively. The displayed tree is the most parsimonious tree, and best supported by pairwise dissimilarity comparison and gene synteny analysis.

Regarding *haoA*, the reconciliation modeled predicted three transfer events (Fig. 6). As for *amoA*, an early transfer from β-AOB to an ancestor of comammox *Nitrospira* is suggested. However, in this case the fact that the similarity between comammox clades is higher than the similarity of each clade compared with the β-AOB genomes may indicate one unique transfer event. Another transfer was inferred from clade B to the ancestor of the clade A genomes A2 and CG24_B. Consistent with this hypothesis, these two genomes contain genes coding for a copper-containing nitrite reductase next to the HAO cluster as is also observed for clade B genomes. Finally, a transfer within β-AOB genomes was detected (Fig. 6).

As comammox clade A and clade B do not form a monophyletic group (Fig. S2), loss events for both *amoA* and *haoA* are expected in some *Nitrospira* spp. phylogenetically placed between these clades such as *N. moscoviensis* and CG24_D. Additionally, the clock-based species-tree estimated the separation between clade A and clade B comammox genomes to have occurred approximately 300 ± 90 MYA, while the split between *Nitrospira* lineage I and II was dated at 375 ± 50 MYA (Fig. S10). As our observations point towards an earlier transfer event for both *amoA* and *haoA*, additional loss events would be required to explain the lack of comammox genes in *Nitrospira* genomes belonging to other lineages than lineage II. One possible explanation for why this might have happened is the tradeoff between rate and yield of ATP production postulated by Costa *et al*. (13). Under this hypothesis, shortening the nitrification pathway leads to an increased growth rate which could be advantageous in some scenarios. Thus the lack of the putatively acquired AMO and HAO-related genes in extant canonical *Nitrospira* spp. could be the result of selection for optimal pathway length.

## Conclusions

In summary, our findings reveal diverse genetic capabilities of the two comammox clades, canonical *Nitrospira* and strict ammonia oxidizers. The decreased presence of NOx detoxification pathways in comparison with canonical AOB together with the diverse genetic capacity for tolerance of low micronutrient concentrations hints at a substrate-limited lifestyle for comammox *Nitrospira*. Additionally, we identified a high probability of transfer events from b-proteobacterial ammonia oxidizers to comammox *Nitrospira* for genes belonging to the ammonium oxidation pathway. Together, these results expand our knowledge of the ecology and evolution of the recently discovered comammox *Nitrospira.*

## Material and Methods

### Recovery of individual genomes

As described in Palomo *et al*.(4), DNA was extracted from triplicate 0.5 g samples taken from two adjacent (ca. 0.5 m apart) locations in a singular RSF (ca. X, Y, Z dimension) and subject to shotgun sequencing to describe the microbial community involved in groundwater purification. The work presented here focuses on the metagenomes from which a *Nitrospira* composite population genome (CG24) was recovered(4). In order to separate the composite population genome into individual genomes, reads from the six metagenomes were independently mapped against subsampled assemblies of the metagenome (ISL_Top2) from which CG24 was extracted (using 5%, 10%, 20%, 30%, 40%, 50%, 60% and 70% of the reads following the strategy described in Hug *et al*. (51)). To recover low abundance genomes, we also co-assembled reads from the three metagenomes where CG24 was most abundant (ISL_Top1, ISL_Top2 and ISL_Top4) and then followed the same strategy as above. Gene calling on the assembled contigs was performed using the metagenome implementation *(‘-p meta*’) of Prodigal v.2.62 (52). A set of 107 HMMs for essential single-copy genes were blasted (BLASTP, E < 1e-5) against the NCBI-nr database with follow-up analysis in MEGAN v.6.5.10 (53) to extract taxonomic assignments from the blast output. With this information, individual genome bins were extracted from each subsampled assembly using mmgenome (54) based on differential coverage (14) (SI Material and Methods and Fig. S1). The quality of recovered genomes was evaluated using coverage plots from mmgenome (54), HMMs for 107 essential single-copy genes and CheckM (55). When the same individual draft genome was recovered from several subassemblies, the one best assembled (based on N50), most complete, and with the lowest contamination was retained. The selected draft genomes were refined using Anvi’o (56) to remove contigs with inconsistent coverage. Assembly quality was improved by alignment against related complete or draft genomes using the Multi-Draft based Scaffolder (MeDuSa) (57) and gaps were closed with GapFiller v.1.10 (58).

### Comparative genome analysis

High quality published *Nitrospira* genomes (more than 90% complete and with less than 5%redundancy) were also included in the comparative genomic analysis. Gene calling on the recovered genome bins and published genomes was performed using Prodigal v.2.62 (52). Annotation was conducted in RAST (59) and protein functional assignments of genes of interest were confirmed using KEGG and blastp. Pangenome analysis was executed using the meta-pangenomic workflow of Anvi’o (56) with default parameters with the exception — maxbit = 0.3. Briefly, blastp was used to calculate similarities between all proteins in each genome. Weak matches between two protein sequences were eliminated using *maxbit heuristic* (60). Finally, the Markov Cluster Algorithm (MCL) (61) was used to generate protein clusters (PCs) from protein similarity search results. PCs were considered part of the core *Nitrospira* genome when present in all *or* all but one of the genomes. The same criterion was used to identify comammox or clade-specific PCs. All comparative genome information was visualized using the program anvi-interactive in Anvi’o (56).

### Sequence dissimilarity and gene synteny analysis

Pairwise amino acid dissimilarities (SI Material and Methods) among the genomes for 18 housekeeping genes, 27 ammonia oxidation-related proteins and 14 syntenic ribosomal protein (Table S1) were calculated using Clustal Omega v.1.2.1 (62). Gene arrangement of ammonium oxidation related genes was visualized using the R package genoPlotR (63).

### Construction of clock-based species-tree

The software BEAST, version 2.4.4, was used to construct a dated species-tree using a relaxed log-normal clock model (64, 65). The input data was a set of 14 ribosomal proteins assumed to be inherited mostly vertically (Table S1). Each protein was aligned individually using the G-INS-i method implemented in the software MAFFT (66), and all alignments were concatenated into a single NEXUS file with one partition per gene. Sequences were assumed to evolve according to the LG substitution model, and to follow a relaxed log-normal clock. Furthermore, each partition was assumed to have its own overall rate, and to have independent gamma-distributed rates over sites. The timetree.org resource (67) together with the substitution rate of 16S rRNA (68) was used as a main starting point for finding calibration information for internal nodes (SI Material and Methods).

### Construction of guest-tree sets

For each investigated gene the protein sequences were aligned using the G-INS-i method in MAFFT (66). The software MrBayes version 3.2.6 (69) was then used to reconstruct phylogenies. MCMC analysis was run for 10 million iterations with a thinning of 10,000 using 2 independent sets of 3 chains and employing the best-fitting substitution model as determined by the program ProtTest3 (70). Convergence was checked visually using the software Tracer, and by checking the average standard deviation of split frequencies for the two posterior tree-distributions. Burn-in (25% of samples) was discarded from the two sets of trees, which were then merged to a single file that therefore ended up containing 1500 tree samples, but corresponding to a much lower number of topologies (individual guest-tree sets contained from 2 to 151 different topologies, with an average of 42). All trees in this file were then midpoint-rooted (SI Material and Methods), thus forming the set of guest-tree topologies that were used for the reconciliation analysis.

### Reconciliation analysis

The analysis of event probabilities was done using the software JPrIME-DLTRS version 0.3.6 (71) (SI Material and Methods). The clock-based tree mentioned above was used as the reference species-tree for all analyses. When the alignment for an investigated gene only contained a subset of the species in the species-tree, we constructed a matching species tree by extracting the relevant sub-tree from the full species tree, updating all branch lengths as necessary to retain patristic distances and ultrametricity. For each gene, the sample file containing 1500 midpoint-rooted trees constructed using MrBayes was used as a guest-tree set from which JPrIME-DLTRS could sample guest-tree topologies. The substitution model for each gene was chosen based on AIC support from the ProtTest3 analysis mentioned above. MCMC was allowed to run for 4 million iterations with a thinning of 200, and running 3 parallel chains for each gene. During the run, samples were also produced for realisations (a realisation, in the terminology of JPrIME-DLTRS, is a reconciliation where the exact time of the event is specified). Convergence was checked by inspecting output using Tracer, and by computing the potential scale-reduction factor (“R-hat”) for model parameters (72). The posterior probability for transfer and duplication events was computed from realization sample files as the fraction of post-burnin samples where the tag “Transfer” or “Duplication” occurred. To investigate which branches the transfer events most likely involved, we used the software ecceTERA version 1.2.4 (73). As a species-tree we again used the fully dated tree constructed from ribosomal proteins. For gene trees we used the midpoint-rooted consensus trees from the JPrIME-DLTRS analysis. We did not consider transfer events to or from extinct or unsampled species (“compute.TD=false”). To account for uncertainty regarding parsimony costs for events, we used strategy “s5” (“pareto.mod=3”) with settings suboptimal.epsilon=1 and real.epsilon=1 (74). For each gene we investigated all reconciliations. The examples presented here were chosen for their consistency with other information (plots of sequence dissimilarities, synteny data and the JPrIME-DLTRS results). The software SylvX version 1.3 (75) was used for visualizing reconciliations computed by ecceTERA.

### Phylogenetic analysis

Phylogenetic trees for AmoA and HaoA were constructed using MrBayes. The substitution model for both data sets was LG with gamma-distributed rates (determined using jmodeltest), using 5 million iterations for 2 sets of 3 chains, with a thinning of 5000 and a burn-in of 25%. Convergence was checked using Tracer and by computing the potential scale reduction factor (ensuring that Rhat was close to 1.0 for all parameters) and the average standard deviation of split frequencies (which was well below 0.01 in both cases).

## Acknowledgements

This research was financially supported by MERMAID (An initial training network funded by the People Programme - Marie Skłodowska-Curie Actions- of the European Union's Seventh Framework Programme FP7/2007-2013/ under REA grant agreement n°607492), and a research grant (Expa-N, 13391) from VILLUM FONDEN.

## Authors Contributions

The study was conceived by AP, BFS and TSP. AP performed the genome binning and comparative genomic analyses. AGP conducted the analysis related to evolution supported by AP. AP led interpretation of the results supported by SJF, AD and AGP. AP drafted the manuscript helped by SJF, AGP, and BSF; all authors contributed to its revision, commented on the manuscript and approved the final submission.

## Additional information

All raw sequence data is available in MG-RAST under accession numbers 4629971.3, 4622590.3, 4629689.3, 4631157.3, 4630162.3 and 4631739.3.

The genome sequences have been deposited at NCBI under the project PRJNA384587, with sequence accession XXX (released after acceptance, please contact for request). The file containing the protein clusters sequences is available on figshare ((http://dx.doi.org/XXX; released after acceptance, please contact for request).

